# Simultaneous analysis of pMHC binding and reactivity unveils virus-specific CD8 T cell immunity to a concise epitope set

**DOI:** 10.1101/2023.11.06.565606

**Authors:** Nikolaj Pagh Kristensen, Edoardo Dionisio, Amalie Kai Bentzen, Tripti Tamhane, Janine Sophie Kemming, Grigorii Nos, Lasse Frank Voss, Ulla Kring Hansen, Georg Michael Lauer, Sine Reker Hadrup

**Affiliations:** Section for Experimental and Translational Immunology, Department of Health Technology, Technical University of Denmark (DTU), Kongens Lyngby, Denmark; Liver Center and Gastrointestinal Division, Department of Medicine, Massachusetts General Hospital and Harvard Medical School, Boston, MA 02114, USA

**Keywords:** Adaptive immunity, virology, CD8 T cells, high-throughput mapping, epitopes

## Abstract

Knowledge of widely recognized T-cell epitopes against common virus infections are vital for immune monitoring and characterization of relevant antigen-specific CD8 T cells and their antigen receptors. We therefore aimed to establish a concise and validated epitope panel for monitoring human virus-specific immunity complete with data on both prevalence of recognition and reactivity in humans. To achieve this, we first establish TCR downregulation, and loss of peptide major histocompatibility (pMHC) multimer-binding, as an early and sensitive marker of T cell reactivity after peptide stimulation. We next applied TCR downregulation in a high-throughput assay by monitoring binding, and loss of binding (i.e. reactivity), to libraries of DNA-barcode labelled pMHC multimers in paired unstimulated/stimulated samples. This novel method allowed us to access T-cell responses in 48 donors towards 929 epitopes recorded in the Immune Epitope Database (IEDB) encompassing 29 virus common infections and 25 different HLA alleles. This yielded a concise panel of 137 virus epitopes, many of which were underrepresented in the public domain, recognized by T cells in peripheral blood. 84% of these epitopes exhibited prevalent reactivity to peptide stimulation, which was associated with effector and long-term memory phenotypes. Conversely, non-reactive responses correlated with naïve and immunosenescence phenotypes. This study represents the largest effort to unbiasedly assess T-cell recognition and reactivity to common virus infections in healthy individuals providing a minimal epitope panel for monitoring adaptive immune responses in humans.

**Significance Statement:** CD8 T-cell epitopes are widely available in public databases yet many are not recognized in the general population. Here we undertook an exhaustive screening process using “state-of-the-art” methods to assess both T-cell recognition and reactivity against common virus infections, which holds significant implications for shaping T-cell immunity and disease protection. We identify 137 commonly recognized epitopes from common virus infections to which T cell responses are expected to occur in human donors. Importantly, several of the verified epitopes were underreported in public databases compared to their observed prevalence of recognition and high cellular frequency making this an important reference dataset and resource for immunologists studying antigen-specific T cells across different immunopathologies and contexts including autoimmunity, infectious disease and cancer immunotherapy.

## Introduction

Antigen recognition by CD8 T cells plays a pivotal role in the adaptive immune response to pathogens, autoantigens, and cancer. Characterization and longitudinal monitoring of the antigen-specific immune response, as well as its underlying antigen-receptor repertoire, is therefore an important and necessary endeavor in humans, particularly in light of the protective and therapeutic potential of such immune responses notably in vaccination^1–3^ and cancer immunotherapy^4–8^, respectively.

Exact epitopes of antigen-specific CD8 T cells can be detected either by binding to cognate peptide major histocompatibility complexes (pMHC) or through a measure of cellular activation in response to antigen stimulation, i.e. “reactivity”^9–14^. However, receptor-ligand binding between a T-cell receptor (TCR) and a pMHC complex is not equivalent to T-cell reactivity to the given epitope^15–17^. This discrepancy may be attributed to various cell intrinsic and extrinsic factors including co-receptor expression^18–20^, TCR affinity to antigen^17,19^, T-cell anergy^18,21^ and chronic antigen stimulation^22^.

To provide a broad, high-quality map of epitopes recognized in context of human virus infections, one therefore needs a sensitive yet high-throughput method with integrated information of both pMHC binding and T-cell reactivity to peptide. This would allow for simultaneous measurement of cohort-wise prevalence of recognition, magnitude (i.e. cellular frequency) and quality of response (i.e. reactivity) integrated into the same epitope map. Current epitope lists, e.g. IEDB^23^, is based on information from very heterogeneous experimental contexts precluding efficient ranking of epitopes according to observed degree of T cell response prevalence and reactivity.

We therefore integrated the high-throughput assessment of TCR-pMHC binding utilizing DNA barcode-labeled pMHC multimers^24^ with determination of T-cell reactivity using synthetic peptide pools^10,25^. Here, T cell specificity is assigned based of the barcode-labeled pMHC multimer binding^24,26,27^. The total number of TCR-pMHC interactions relevant for a given T cell population can furthermore be assessed through the embedded unique molecular identifiers (UMIs)^28^. A reduction in TCR surface expression, due to activation-induced downregulation^29–33^, will therefore lead to a reduced pMHC surface density and hence reduced barcode-associated UMI counts for a given specificity. As such, T-cell reactivity, in form of TCR downregulation following antigen exposure, can be assigned for each specific T-cell population, by measuring the relative loss of signal for a specific barcode associated with its unique pMHC multimer. We then tested epitope-binding and reactivity of 10,365 pMHC interactions across 48 donors allowing us to define 137 high-confidence epitopes across common virus infections along with 2 epitopes from tumor-associated antigens, that form a hierarchy of functional epitope-recognition in blood donors and a reference map complete with expected values for prevalence of recognition and reactivity. Moreover, we demonstrate here that non-functional T cell characteristics are often linked to T cell phenotypes corresponding either to lack to antigen exposure (i.e. naïve T cells) or immunosenescence (CD57^hi^ Temra cells) and that certain epitope frequently drives such non-functional T cell responses.

## Results

### Diminished pMHC-multimer binding is an early and sensitive marker of T-cell reactivity

TCR downregulation and the associated decreased pMHC multimer staining of recently activated antigen-specific CD8 T cells has been documented for both high and low avidity ligands in the OT-I and 2C TCR transgenic systems^32^. Similar observations have been made *ex vivo* in a human setting using selected epitopes derived from human cytomegalovirus (CMV) and influenza virus (FLU)-antigens^33^. We explored this phenomenon in detail for a variety of well-described viral epitopes using human PBMCs. We observed strongly decreased median fluorescent intensities (MFI), and decreased frequencies of pMHC multimer positive T cell populations following 24-hour peptide stimulation across all virus-derived epitopes tested (Fig. 1a-c, Supplementary Fig. 1a). This coincided with the upregulation of canonical activation markers CD69+ and CD137+ (AIMs)^10,12^ (Fig. 1a, d). Furthermore, dilution of the stimulating peptide showed a clear dose dependency (Fig. 1e, Supplementary Fig. 1b-c). Finally, we performed a time series to map the temporal kinetics of the effects of peptide stimulation in the context of different antigen-specific populations from the same donor. We observed a decrease in pMHC multimer binding specific to the targeted CD8 T-cell response as early as 3 hours post stimulation, with the strongest effect at 24-48 hours. We moreover found no evidence of renewed capacity to bind pMHC multimers at 48 hours post stimulation (Fig. 1f, Supplementary Fig. 1d-g). CD137+ kinetics were largely similar to previously published reports (Supplementary Fig. 1h)^12^, although we observed a tendency for marked background staining of anti-CD137 particularly at early time points, (Supplementary Fig. 1i). In summary, we confirm reduction in pMHC multimer binding to T cells as a specific and sensitive readout for productive TCR engagement in vitro.

**Figure 1.**
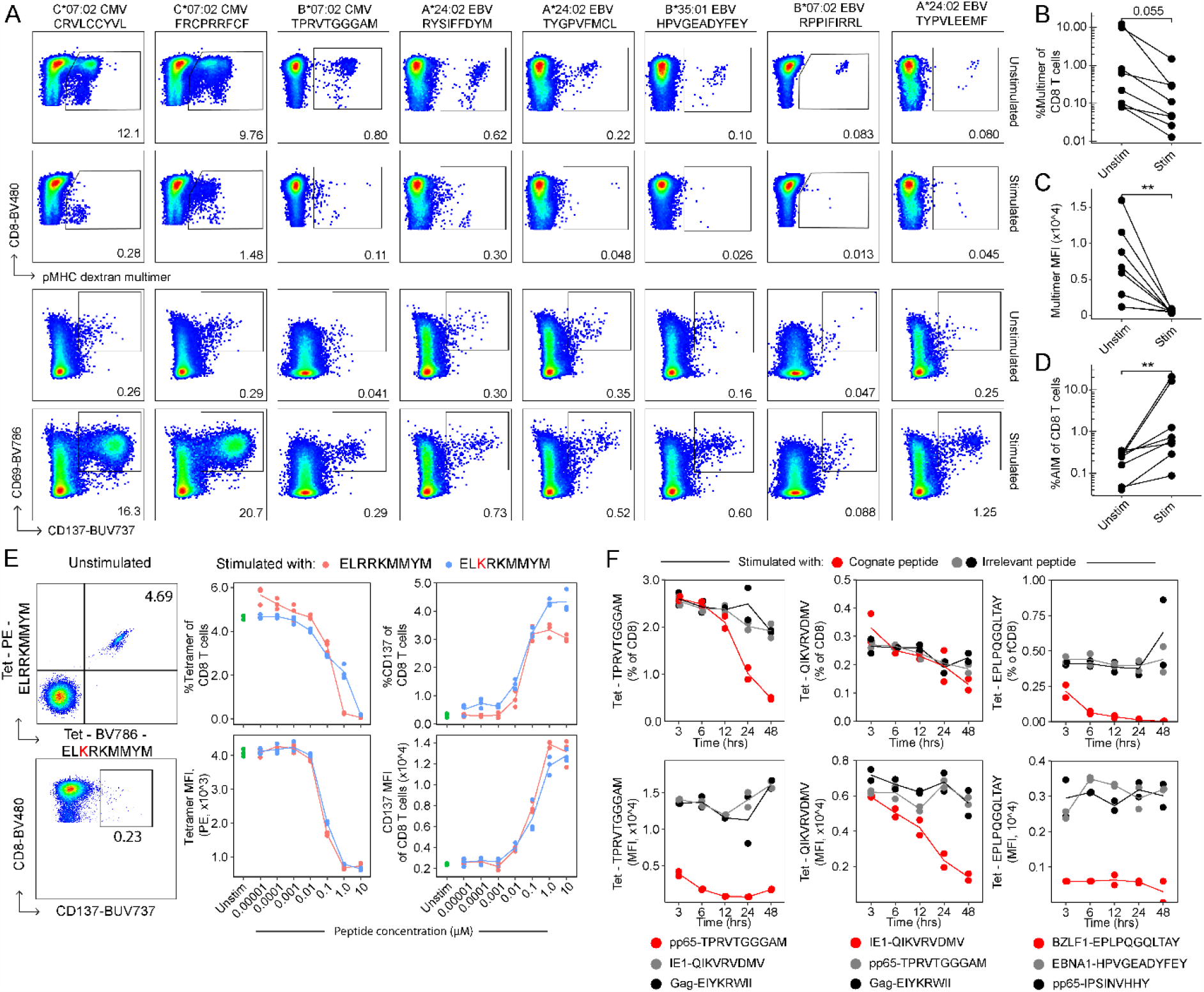
Diminished intensity of pMHC multimers is an early and sensitive marker of T cell activation. (A) Loss of pMHC multimer-staining following peptide-based stimulation of antigen-specific CD8 T cells and concomitant increase in frequency of CD69+CD137+ CD8 T cells. PBMCs were stimulated with 1 μM peptide for 24 hours. Peptide and pMHC multimer identity indicated in the header for each column. Unstimulated controls were treated with equimolar DMSO. Samples were pregated for live CD14-CD19-CD3+CD4-CD8+ lymphocytes. (B-D) Statistics pertaining to A. (B) Multimer frequency of CD8 T cells, (C) multimer MFI, (D) AIM frequency i.e. frequency of CD69+CD137+ of CD8 T cells. (E) Limiting dilution of two 24-hour stimulation series in triplicates. Unstimulated controls were treated with 10 μM DMSO. Tetramer+ cells were pregated on live CD3+CD4-CD8+ lymphocytes. Tetramer MFI and frequency were measured. Tetramer MFI for ELKRKMMYM-BV786 is shown in Supplemental Figure 1c. (F). Time series spanning 3-48 hrs of stimulation with 1 μM peptide in duplicates. Each time series was stimulated with one of three peptides, either one of two irrelevant peptides or a cognate peptide, and subsequently stained with all three tetramers. Tetramer+ cells were pregated on live CD3+CD4-CD8+ lymphocytes. Tetramer MFI and frequency were measured. Tetramer-null controls were used as gating controls, see Supplemental Figure 1d. Representative flow plots are shown in Supplemental Figure 1e-f. Two additional positive peptides are presented in Supplemental Figure 1g. p-values were calculated using a paired Wilcoxon test. **, *p < 0*.*01*. MFI, median fluorescent intensity. AIM, activation-induced markers.

### Peptide stimulation correlates with loss of pMHC multimer binding measured by DNA barcode counts in antigen-specific CD8 T cell populations with negligible bystander effects

To scale our experimental workflow of assessing both T cell epitope binding and reactivity simultaneously (Fig. 2a) to hundreds of epitopes using DNA barcode-labeled multimers, we selected 945 epitopes from 29 different common viruses (Supplementary Table 1) restricted to 25 HLA alleles from IEDB (selection details in methods). Epitopes were selected to represent viruses with frequent exposure in healthy populations. The epitope library was characterized by a predominance of herpesvirus epitopes as well as HLA-A*02:01 restricted peptides (Fig. 2b, Supplementary Fig. 2a). As a reference, the library also included two epitopes derived from well-known tumor-associated self-antigens; MART1_26-35(Leu27)_ ELAGIGILTV and NY-ESO_157-165_ SLLMWITQV restricted to HLA-A*02:01^34,35^.

**Figure 2.**
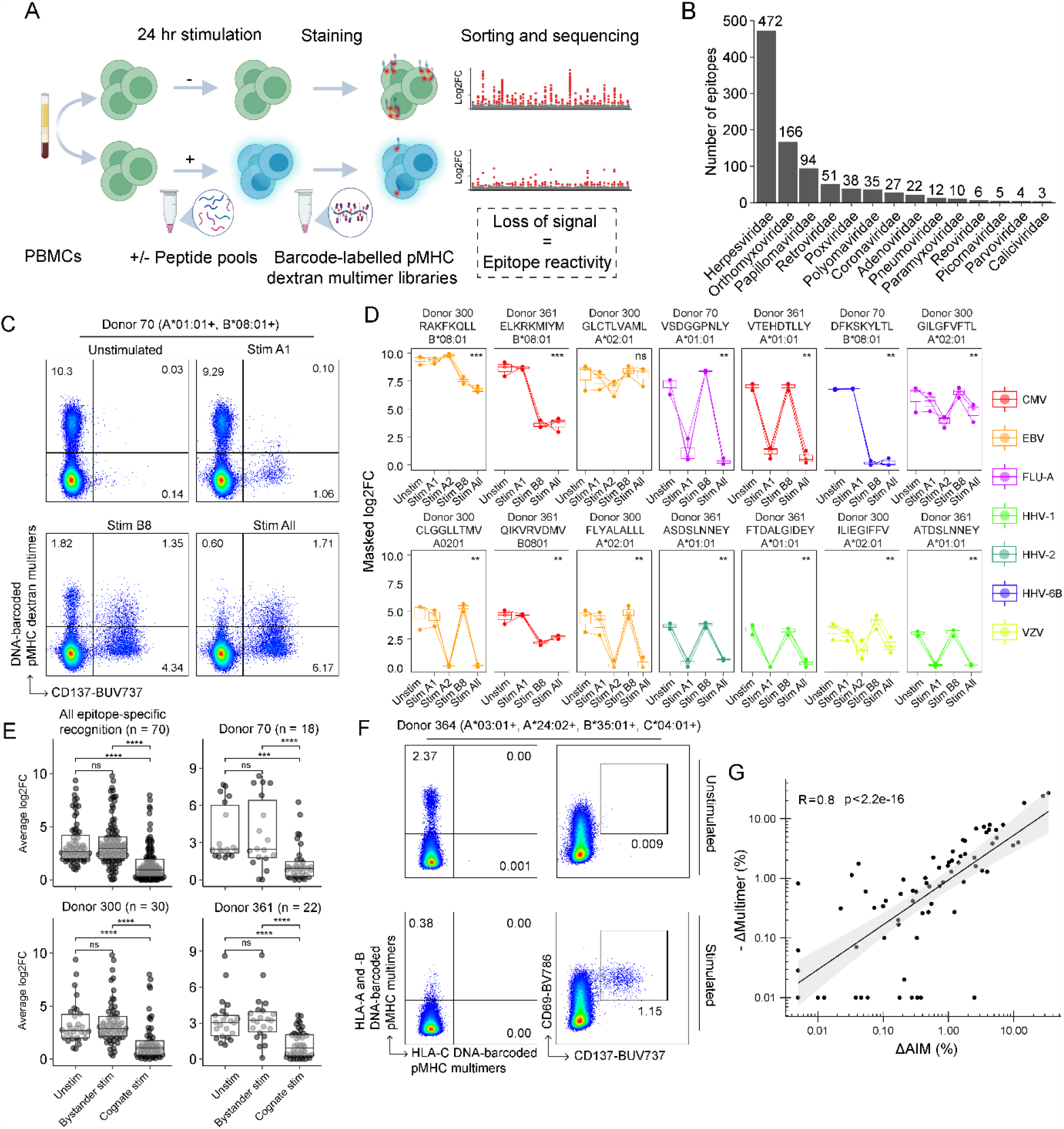
Multiplexing pMHC multimers using DNA-barcodes allow monitoring of specific activation following peptide pool-based stimulation. (A) Biorender®-generated flow chart describing the general work flow including sample input and sequencing output. (B) Multimer-library size containing epitopes from distinct virus families selected from IEDB (see methods). (C-D) Assessment of multimer-binding following stimulation in three donors using technical triplicates. (C) Representative cell sorting of antigen-specific CD8 T cells using barcode-labelled pMHC multimers following different pool-based stimulations for 24 hours. Cells from all upper quadrant gates were sorted. Stim A1, Stim B8 and Stim All refer to pool-based stimulations with 36 A*01:01-restricted peptides, 50 B*08:01-restricted peptides and the collective pool of 86 peptides, respectively. All conditions were recorded in triplicates. (D) Enrichment scores for select multimer-specific CD8 T cells across three donors and 3-4 stimulatory settings. Stim A2 refers to the pool-based stimulations with 96 A*02:01-restricted peptides. Samples were pregated for live CD14-CD19-CD3+CD4-CD8+ lymphocytes. See additional populations in Supplemental Figure 2f-g. (E) In total, 70 virus-specific populations were discovered and grouped according to whether the respective stimulatory setting included the peptide used for multimer-generation (cognate stim) or not (bystander stim). (F) PBMCs from 48 donors were stimulated individually with pools ranging 1-6 donor-derived HLA alleles. Multimer positive CD8 T cells were then sorted from both unstimulated and stimulated cultures for barcode-sequencing. Samples were pregated on live CD14-CD19-CD3+CD4-CD8+ lymphocytes. Additional representative flow cytometry is available in Supplemental Figure 2h. (G) Correlation between observed change in multimer frequency (Δmultimer(%)) and change in CD69+CD137+ frequency (ΔAIM(%)) upon stimulation across all 48 donors. Correlation analysis was performed using Spearman’s correlation. Post hoc analysis was performed using Dunn’s test and p-values were adjusted using Benjamini-Hochberg. Unpaired Wilcoxon test between grouped unstimulated/bystander stimulated samples and cognate stimulated samples was performed in D. Boxplot bounds show the 25^th^ and 75^th^ percentiles along with the median. Upper and lower whiskers show the range of data unless data points are lower/larger than 1.5xIQR. *, *p < 0*.*05*. **, *p < 0*.*01*. ***, *p < 0*.*001*. ****, *p < 0*.*0001*.

We verified that activation-induced TCR downregulation and reduction in pMHC multimer staining intensity was (1) measurable by DNA barcodes used to tag individual pMHC multimers, and (2) unaffected by activation of other antigen-reactive CD8 T cells (i.e. bystander activation). To do this, we included a wide panel of DNA barcode-labelled pMHC multimers encompassing 36, 96 and 59 virus epitopes restricted to HLA-A*01:01, A*02:01 and B*08:01, respectively. We split PBMCs from 3 healthy donors into unstimulated, partially stimulated and fully stimulated short-term cultures, and stained each culture in triplicates with donor-specific, HLA-matching pools of barcoded multimers. Amplicon sequencing of the pMHC multimer associated barcodes was subsequently performed on multimer+ CD8 T cells, and individual enrichment scores for TCR:pMHC binding calculated based on such barcodes. As expected, we observed loss of fluorescent pMHC multimer binding when using HLA-A*01:01, A*02:01 or B*08:01-restricted peptide pools (Fig. 2c, Supplementary Fig. 2b) and a corresponding increase of CD137 expression (Supplementary Fig. 2c). Furthermore, we observed that cognate peptide stimulation resulted in decreased enrichment scores (masked log2FC) for specific pMHC multimer binding, while T cell populations specific for other epitopes in the same sample were unaffected by such stimuli (Fig. 2d-e). In total, 70 epitope-specificities were assessed from 3 donors, and the majority of epitopes (54 of 70) were significantly affected by peptide stimulation (Fig. 2d, Supplemental Fig. 2d-e). Next, we scaled our methodology to the final panel of 929 virus epitopes (see methods for excluded epitopes), and evaluated T cell responses in 48 donors (27 healthy Danish blood donors and 21 American donors diagnosed with acute or chronic viral hepatitis). Peptides were grouped and pooled according to HLA-restrictions as reported in IEDB and recombined on the day of stimulation into donor-specific peptide pools covering up to 6 HLA class I alleles for any given donor. DNA barcode-labeled MHC multimers were also grouped according to HLA and combined into donor-specific multimer pools. All multimer-binding T cells were then sorted to determine donor-specific T cell responses (Fig. 2f, Supplementary Fig. 2f). While donor-to-donor variations was observed, the increase of AIM+ CD8 T cells in stimulated samples corresponded to the observed decrease of multimer-specific CD8 T cells (*R* = 0.8, *p* < 2.2e-16) (Fig. 2g). Furthermore, the general increase in CD137 MFI values also corresponded to decreased multimer fluorescent intensities (R = 0.6, *p* = 5.7e-4) (Supplementary Fig. 2g).

Overall, these data establish that an assay combining pMHC multimer staining with peptide stimulation can be used to simultaneously test hundreds of distinct T cell specificities with DNA-barcoded pMHC multimer libraries and determine T cell antigen reactivity with negligible bystander effects.

### Evaluation of pMHC binding and antigen reactivity for large epitope libraries identifies a hierarchy of antigen-specific CD8 T cells with different reactivity levels

Next, we assessed TCR downregulation for individual antigen-specific CD8 T cell responses across our cohort, by accounting for decreased UMI counts for any barcode. We studied a total of 10,365 pMHC interactions and identified 647 epitope-specific CD8 T cell populations in unstimulated samples. We estimated the underlying frequency of epitope-specific CD8 T cells by weighting the total multimer+ population of CD8 T cells observed by flow cytometry according to the fraction of mapped UMIs pertaining to the a given epitope^4,24^, which resulted in an estimated frequency range of 0.0005%-18.7% of total CD8 T cells in unstimulated samples. All identified responses across the total cohort are depicted in Fig. 3a (upper panel, unstimulated sample), each dot representing one multimer-specific population in a given donor based on significantly enriched pMHC barcodes. Parallel assays after epitope peptide pool stimulations (Fig. 3a, bottom panel) revealed a concomitant partial binding loss across a wide range of T cell specificities targeting different epitopes and viruses. Δlog2FC values were subsequently calculated comparing paired multimer data from unstimulated and stimulated samples, where negative Δlog2FC generally represents decreased multimer-binding (i.e. reactivity) and positive values indicate unchanged binding. T cell populations assigned with antigen reactivity (Δlog2FC<0) are circled, while non-reactive populations (Δlog2FC>0) are triangled (Fig. 3a). In total, 564 of the 647 identified epitope-specific populations were assigned with a net negative Δlog2FC after stimulation, whereas 83 responses were assigned with positive or unchanged Δlog2FC values (Fig. 3b).

**Figure 3.**
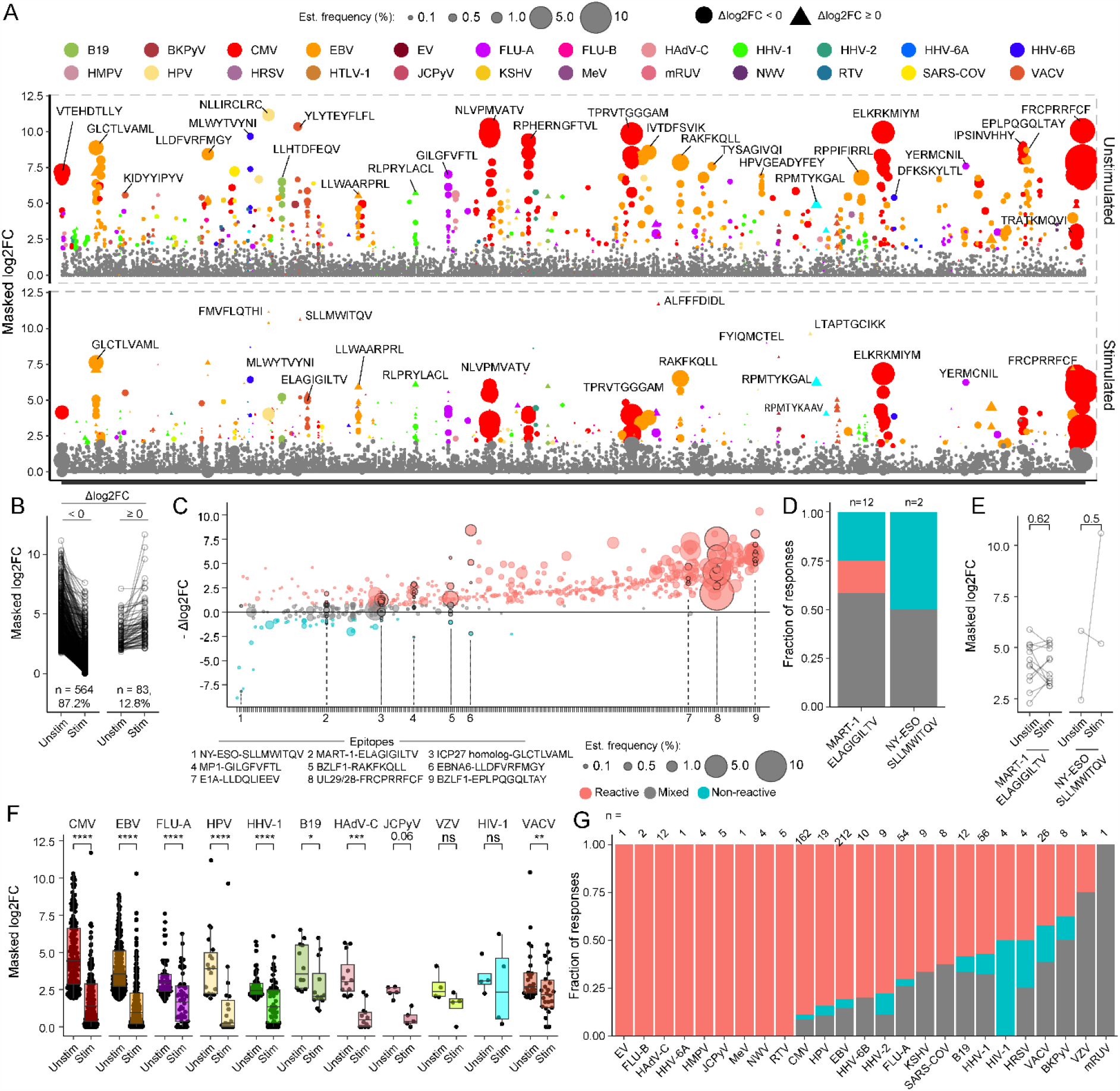
Simultaneous evaluation of epitope binding and reactivity for large epitope libraries captures a hierarchy of antigen-specific CD8 T cells with heterogeneous antigen reactivity. (A) Epitope recognition for all 48 donors and paired conditions (unstimulated PBMCs in top panel vs stimulated PBMCs in bottom panel). Significantly enriched pMHC multimers are colored corresponding to their virus of origin. T cell populations with Δlog2FC<0 are circled, and T cell populations with Δlog2FC>0 are triangled. Size of the dots denotes estimated frequency of the underlying epitope-specific CD8 T cells. (B) Barcode enrichment scores comparing unstimulated samples to stimulation samples. (C) Epitope-recognition including underlying estimated frequency (dot size) depicted according to the observed reactivity score (-Δlog2FC). Each epitope and corresponding epitope-specific populations are ranked on the horizontal axis according to average Δlog2FC and grouped in three bins: -Δlog2FC > 0.75 (“Reactive”), −0.75 < - Δlog2FC < 0.75 (“Mixed”), -Δlog2FC < −0.75 (“Non-reactive”). Notable responses are highlighted with black borders and their identities specified on the bottom axis. (D) Proprotion of Reactive, Mixed and Non-reactive epitope-recognition among tumor-specific CD8 T cells recognizing MART-1-ELAGIGILTV and NY-ESO-SLLMWITQV restricted to HLA-A*02:01. Numbers above each column denote the total number of independent responses discovered for each epitope-specificity. Reactive, Mixed and Non-reactive categories of responses, annotated as in C. (E) Masked log2FC score for each multimer and DNA-barcode associated with tumor-shared antigens MART-1-ELAGIGILTV and NY-ESO-SLLMWITQV comparing unstimulated samples to stimulation samples. (F) Pairwise comparison of multimer enrichment scores following stimulation for representative virus origins, see Supplemental Figure 3d for all virus groups. (G) Proprotion of Reactive, Mixed and Non-reactive epitope-recognition among virus-specific CD8 T cells across all donors. Numbers above each column denote the total number of independent responses discovered for each virus origin. Only multimers recognized in at least two independent donors were included. Masked log2FC values set negative barcode enrichment scores (log2FC) to 0. *p*-values in in E and F were calculated using paired Wilcoxon test. Boxplot bounds show the 25^th^ and 75^th^ percentiles along with the median. Upper and lower whiskers show the range of data unless data points are lower/larger than 1.5xIQR. *, *p < 0*.*05*. **, *p < 0*.*01*. ***, *p < 0*.*001*. ****, *p < 0*.*0001*.

In order to generate discrete categories of TCR reactivity that take into account areas of technical uncertainty – we regarded -Δlog2FC > 0.75 as “reactive” epitope recognition; −0.75 < −Δlog2FC < 0.75 as a “mixed” population of reactive and non-reactive epitope-recognition; and -Δlog2FC > 0.75 as “non-reactive” epitope recognition. This allowed us to visualize and rank epitope-specific recognition according to measured TCR downregulation and loss of pMHC multimer binding across multiple independent donors (Fig. 3c). Examples of highly reactive populations (indicated by numbers on Figure 3C) included B*35:01-restricted BZLF1-EPLPQGQLTAY from Epstein-Barr virus (EBV, mean est. frequency: 0.26%), A*02:01-restricted E1A-LLDQLIEEV from HAdV-C (mean est. frequency: 0.23%) and C*07:02-restricted UL29/28-FRCPRRFCF from CMV (mean est. frequency: 7.16%). Examples of non-reactive populations featured recognition of A*02:01-restricted NY-ESO-SLLMWITQV (mean est. frequency: 0.02%) and to some extent MART1-ELAGIGILTV (mean est. frequency: 0.11%), which were analyzed in 2 and 12 independent donors, respectively (Fig. 3d-e).

To explore reactivity hierarchies between T cell responses targeting different viruses, we grouped T cell recognition of epitopes derived from the same virus together and compared their overall enrichment scores as well as their frequency of reactive, mixed and non-reactive response patterns. (Fig. 3f-g, Supplementary Fig. 3a). A significant reduction in barcode enrichment scores were found for a range of viruses including EBV, CMV, FLU-A, human papillomavirus (HPV), herpes simplex virus 1 (HHV-1), B19 parvovirus (B19), human mastadenovirus C (HAdV-C), JC polyomavirus (JCPyV, p = 0.06), vaccinia virus (VACV), Kaposi’s sarcoma-associated herpesvirus (KSHV), human metapneumovirus (HMPV), human herpesvirus 6B (HHV-6B), severe acute respiratory syndrome coronavirus (SARS-COV), and rotavirus A (RTV, p = 0.06) (Supplementary Fig. 3a). However, when taking into account donor-specific heterogeneity, we found that VACV, HHV-1, BKPyV and HIV-1 in particular showed a bimodal distribution, with reactivity assigned only in approximately half of the donors (Fig. 3g).

In summary, our methodology enabled large-scale discovery of virus-specific CD8 T cells across a cohort of PBMC donors, revealing a reactivity hierarchy between virus infections.

### A reference map of high-confidence virus-derived CD8 T-cell epitopes and their associated antigen-reactivity

Ranking epitope-recognition by measured antigen-reactivity adds important additional information when generating reference maps of epitopes recognized in cohorts of interest. To derive a reference map of high-confidence CD8 T-cell virus-epitopes from common virus infections in our donors, we selected all epitopes recognized in at least two independent donors and ranked them according to proportion of reactive, mixed and non-reactive epitope-specific CD8 T cell populations among the detected responses (Fig. 4a). In total, we report 137 virus epitopes out of which 84% (115 epitopes) featured in ≥50% of cases reactive T cell responses. 16 epitopes were dominated by T cell responses with mixed reactivity. Finally, we observed that EBNA4-RAKFKQLL, LMP2-RRRWRRLTV, EBNA-1-FVYGGSKTSL, ICP22-APRIGGRRA, LMP2-PYLFWLAAI and D1-RPSTRNFFEL were more heterogeneous, with no single of the three categories exceeding 50%. We present data on cohort-wide antigen-reactivity in Fig. 4a and Supplementary Table 2 along with the prevalence of CD8 T-cell recognition across our cohort, the average population frequency observed in peripheral blood, the number of IEDB reports associated to a given epitope, the average ‘reactivity score’ Δlog2FC, and the predicted MHC binding affinity for each epitope (netMHCpan v. 4.1)^36^. We observed a similar frequency of virus-specific recognition between American and Danish donors, although Danish donors tended to have increased frequency of B19, FLU-A and HHV-1-specific CD8 T cells (Supplementary Fig. 3b-c). CMV and EBV-specific CD8 T cell populations were typically larger as indicated by median frequencies of 0.141% and 0.09%, respectively, compared to FLU-A, HHV-1 and KSHV-specific CD8 T-cell populations with median frequencies at 0.03%, 0.04% and 0.03% (Fig. 4b), respectively, supporting previous findings^37^. CMV-specific CD8 T cells were also detected at a statistically higher frequencies than HPV and polyoma-specific CD8 T cells, although the latter entailed too few observations to adequately estimate the *p*-value. Interestingly, epitope-specific CD8 T cells targeting VACV, B19, HadV-C and HHV-6B were present at similar median frequencies as CMV and EBV at 0.07%, 0.09%, 0.1% and 0.13%, respectively (Fig. 4b). We also summarized the number of high-confidence epitopes found for each HLA allele. As expected, due to the large number of IEDB-derived candidate epitopes restricted by HLA A*02:01, most of the identified 137 virus epitopes in our screening were restricted by this allele (Fig. 4c, Supplementary Fig. 2a). These data establish a reference map of virus-specific CD8 T cells based on a large library of viral antigens across multiple viruses and HLA alleles.

**Figure 4.**
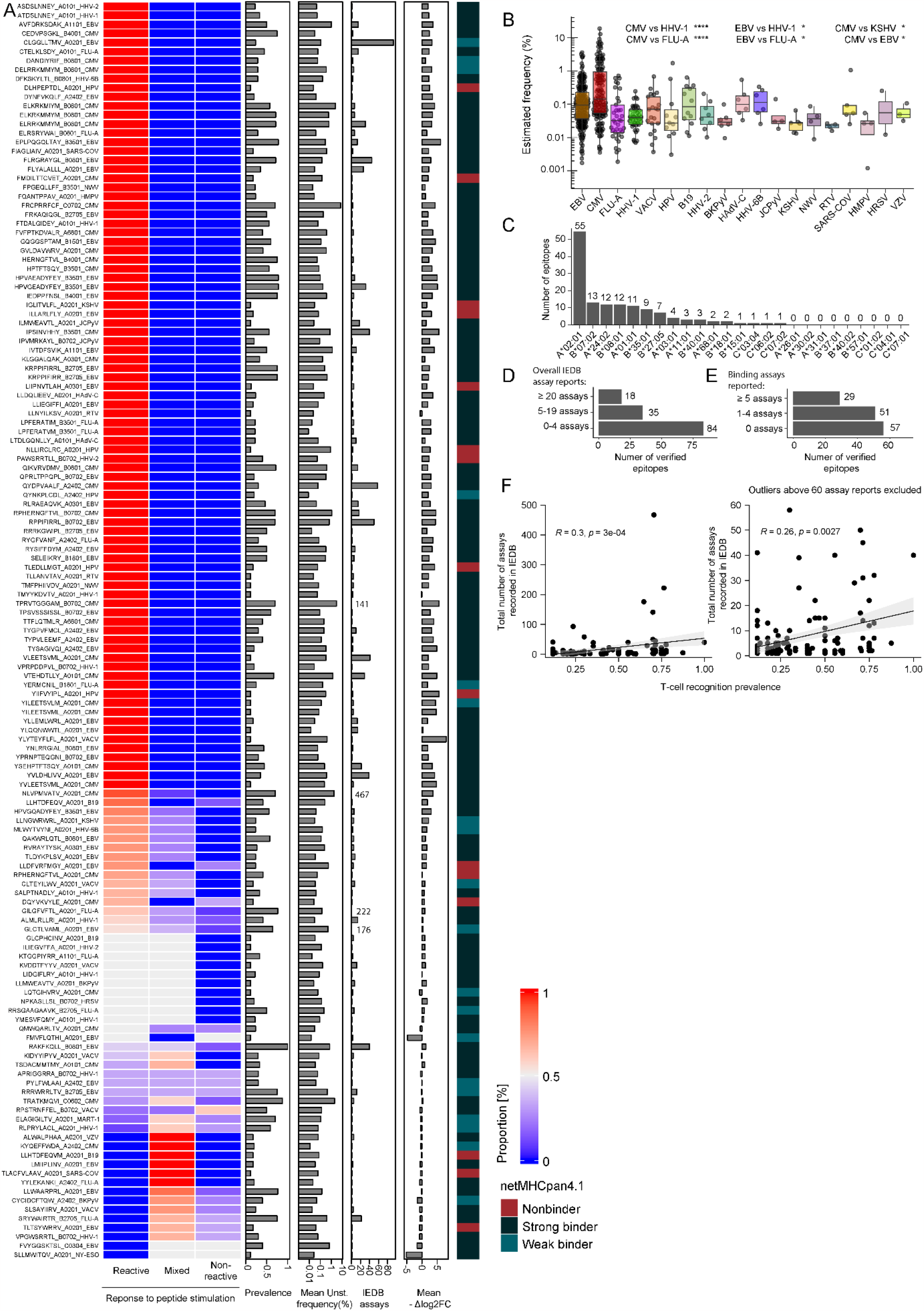
Common virus epitopes associated with reactivity and non-reactivity. (A) Heatmap denoting proportion of reactive, mixed or non-reactive epitope-specific CD8 T cell populations. Only T cell epitopes recognized by at least two independent donors are depicted. Additional panels denote prevalence of detection in our cohort, the average response frequency observed in unstimulated control samples, the number of previous positive IEDB assay reports associated with epitope recognition and the mean reactivity score (Δlog2FC), respectively. The rightmost colored boxes denotes predicted pMHC-binding calculated by netMHCpan4.1 (Reynisson et al., 2021) for each epitope. (B) Estimated frequency in unstimulated samples for all responses detected grouped by virus origin. (C) Number of commonly recognized epitopes per HLA. (D) Number of commonly recognized epitopes grouped in three bins according to number of reported assays in IEDB at the time of epitope selection. Structural assays such as X-ray crystallography were excluded from this count (see materials and methods). (E) Number of epitopes associated with minimal reactivity grouped in three bins according to number of reported qualitative binding assays in IEDB including pMHC multimers. (F) Correlation between observed prevalence and IEDB assay reports for all 109 epitopes associated with common reactivity. Left: All data. Right: Outliers with extreme number of reports were exclude (above 65 assays). All metrics from IEDB stem from the original database search Oct. 2020. Boxplot bounds show the 25^th^ and 75^th^ percentiles along with the median. Upper and lower whiskers show the range of data unless data points are lower/larger than 1.5xIQR. Correlations and associated p-values were calculated using Spearman correlation. p-values were calculated in B using Dunn’s test and adjusted using the Benjamini-Hochberg method. ^*^, *p < 0*.*05*. ^**^, *p < 0*.*01*. ^***^, *p < 0*.*001*. ^****^, *p < 0*.*0001*.

Among the 137 high-confidence epitopes, for which T cell recognition was detected, several were minimally described in IEDB (Fig. 4d-e). This was furthermore visible from the numerically significant yet weak correlation between IEDB assay reports and observed prevalence of reactivity (*R* = 0.26-0.30, Fig. 4f).

In summary, we report recognition of 137 common virus epitopes along with corresponding antigen-reactivity, prevalence, and mean frequency of T cell recognition, by comparing all epitopes in a single experimental framework.

### Differences in antigen-reactivity reflect co-receptor expression and effector differentiation

To investigate phenotypic characteristics of reactive and non-reactive responses, we selected 12 donors for verification using conventional pMHC tetramers and immunophenotyping including specificities from CMV, EBV, FLU-A, B19, VZV, HAdV-C, HHV-1, HHV-6B and MART-1. We ranked each donor-derived T cell response according to reactive, mixed and non-reactive categories introduced in Fig. 3. Up to six specificities were encoded per staining reaction using up to two fluorescent tetramers (Fig. 5a). We found that virus-specific CD8 T cell frequencies assessed using conventional tetramers closely matched with the estimated frequencies acquired from the initial barcoded pMHC multimer screens (R=0.7, p=3.5e-10) (Fig. 5b). However, we observed pronounced phenotypic heterogeneity between epitope-specific T cells recognizing different virus antigens of interest (Fig. 5c-d, Supplementary Fig. 4a-b). Epitope-specific CD8 T cells recognizing CMV and B19 infections were for example dominated by effector memory T cells (Tem, CCR7-CD45RA-), whereas epitope-specific CD8 T cells recognizing FLU-A, HAdV-C and HHV-1 exhibited central memory (Tcm, CCR7+CD45RA-) and stem-like memory (Tscm, CCR7+CD45RA+CD95+) phenotypes (Fig. 5d). Recognition of CMV epitopes was additionally associated with expression of GzmB (Fig. 5d).

**Figure 5.**
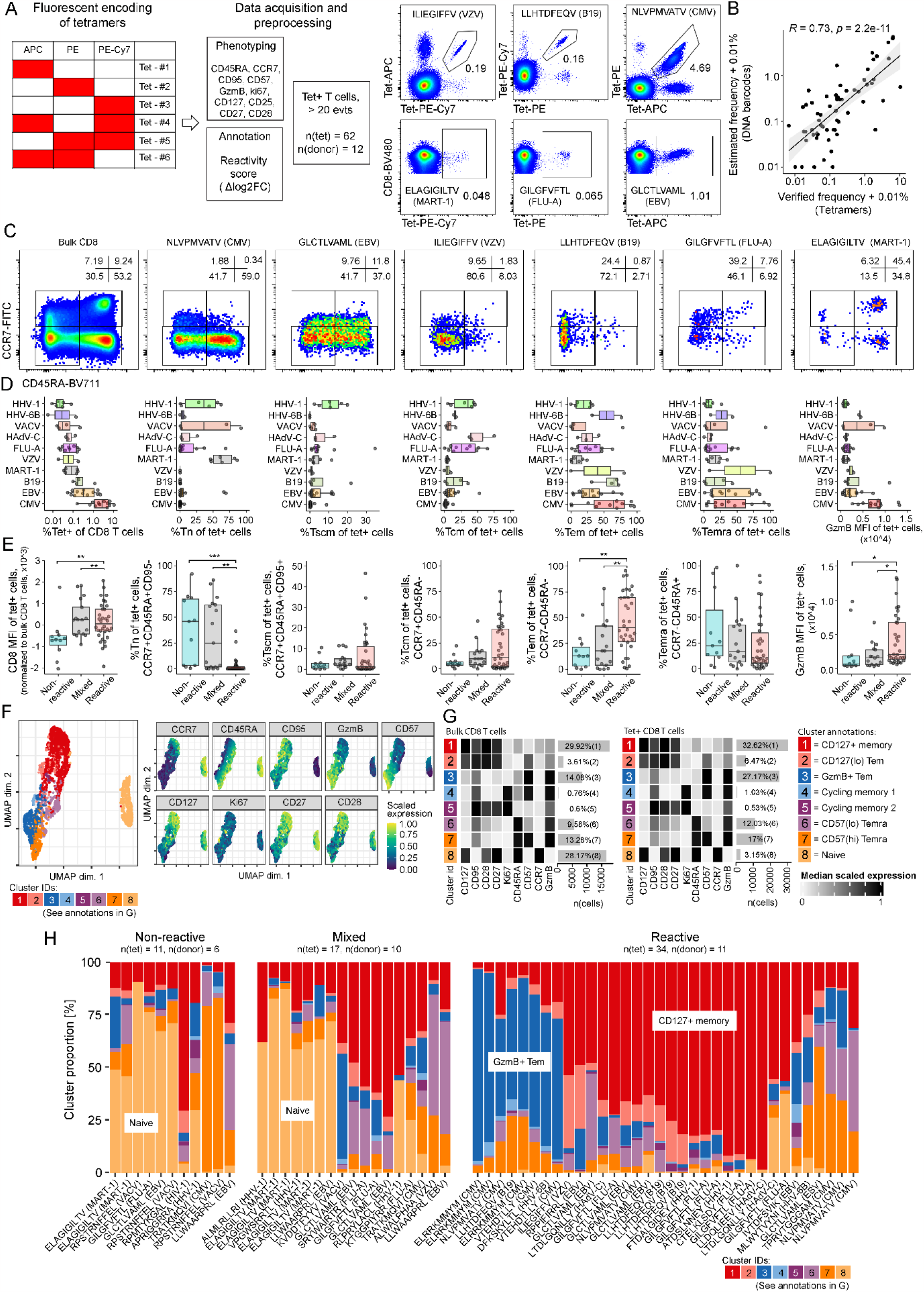
Differences in antigen-reactivity reflect co-receptor expression and effector differentiation. (A) Combinatorial encoding of fluorescent pMHC-tetramers, experimental setup and data processing for verification of 62 different epitope-specific CD8 T cell populations. Antigen-specific CD8 T cells for a given fluorophore combination were pregated on CD3+CD4-CD8+ T cells negative for irrelevant tetramer fluorophores. Representative tetramer gating is shown from donor 310. (B) Correlation (Spearmann) between observed frequency using DNA-barcodes (see methods) and verified frequency using fluorescent tetramers. (C) Markers of memory and effector differentiation CCR7 and CD45RA for bulk and tetramer positive CD8 T cells of donor 310. (D) Summary on tetramer frequency, memory phenotypes and GzmB expression measured by flow cytometry for antigen-specific T cells split by virus origin. (E) Quantification of CD8 MFI, Tn, Tscm, Tcm, Tem, Temra and GzmB MFI for tet+ CD8 T cells grouped by their observed reactivity from initial stimulation and multimer screening. (F) Dimensionality reduction of cytometry data using flowSOM and UMAP. Maximum 5000 tetramer+ from each population of tetramer+ CD8 T cells, and maximum 5000 bulk CD8 T cells were included from each donor. Maximum 50 random cells from each of tetramer+ and bulk CD8 T cell data files were used for the generation of the UMAP. Marker genes used for FlowSOM clustering are shown along their median scaled expression value. Cluster annotation are shown in G. (G) Heatmaps representing 8 distinct clusters. Eight clusters were used to minimally cover Tn, Tem,Temra along with cycling cells and CD127+ memory cells (see methods). (H) Proportion of all 8 clusters represented among individual epitope-specific CD8 T cells grouped by response category (Non-reactive, Mixed and Reactive). All tet+ populations (n = 62) were included. *p*-values were calculated using Dunn’s test and adjusted using the Benjamini-Hochberg method. ^*^, *p < 0*.*05*. ^**^, *p < 0*.*01*. ^***^, *p < 0*.*001*. ^****^, *p < 0*.*0001*. Tet, tetramer.

When grouped into non-reactive, mixed and reactive responses, we observed that CD8 co-receptor density was strongly reduced for the majority of non-reactive epitope-specific CD8 T cell responses (Fig. 5e). Additionally, we observed a high propensity for naïve phenotypes (Tn, CCR7+CD45RA+CD95-) among non-reactive epitope-specific CD8 T cells (Fig. 5e). This is consistent with prior literature showing that naïve CD8 T cells can exhibit reduced TCR sensitivity due to low surface expression of the CD8 co-receptor^18^. The lack of reactivity among naïve CD8 T cells also aligns with previous description of anergic MART-1-ELAGIGILTV specific T cells in healthy (non-vitiligo, non-melanoma) donors^21,38^. We additionally observed increased proportion of Tem cells among reactive epitope-specific CD8 T cells and a reduced proportion among mixed and non-reactive epitope-specific CD8 T cells (Fig. 5e). Tcm cells and effector memory re-expression CD45RA (Temra, CCR7-CD45RA+) populations were more evenly distributed among the three categories of antigen reactivity.

We performed FlowSOM clustering to further explore phenotypic subsets within non-reactive, mixed, and reactive epitope-specific CD8 T cells. We then generated a two dimensional representation of phenotypic markers using Uniform Manifold Approximation and Projection (UMAP)(Fig. 5f). Cluster 1 and 2 were identified as memory CD8 T cells with varying degree of CD127 expression, cluster 3 consisted of GzmB+ Tem cells, cluster 4-5 consisted of Ki67+ cycling cells, cluster 6-7 consisted of Temra cells and cluster 8 consisted of naïve CD8 T cells which were clearly separated from memory CD8 T cells (Fig. 5g). Non-reactive epitope-specific CD8 T cells were again dominated by naïve phenotypes (Fig. 5h). However, two non-reactive epitope-specific CD8 T cells also exhibited high propensity of CD57(hi) Temra phenotypes without expression of CD27 and CD28 (Fig 5h). This phenotype is consistent with senescent T cell populations which accumulate in aged individuals^39^. FlowSOM generated clusters that enriched among reactive CD8 T cells included GzmB+ Tems as well CD127+ memory cells (Fig. 5h). Interestingly, GzmB and CD127 were seemingly counter-expressed in our analysis (Fig. 5g). GzmB+ Tem and CD127+ memory cells therefore likely represent two independent subsets with high TCR sensitivity for antigen stimulation.

We additionally examined selected T cell populations based on single-cell transcriptomics. T cell populations were selected based on pMHC multimer binding, sorted and analyzed through the 10x Chromium platform. We assigned both pMHC specificity, TCR usage and transcriptomic features for all captured CD8 T cells. We evaluated 11 different T cell populations, and observed that high-frequency T cells with lower mean reactivity (BZLF1-RAKFKQLL) demonstrated distinct phenotype characteristics (*GZMK, HLA-DR, CD74, KLRB1, CRTAM*), while the most reactive T cell populations displayed *GNLY and GZMB* expression (Supplementary Fig. 4).

Taken together, we conclude that TCR downregulation and sensitivity to antigen-stimulation is associated with distinct T-cell phenotypes. Naïve as well as senescent phenotypes associate with reduced TCR sensitivity to antigen-stimulation, whereas GzmB+ Tem cells and CD127+ memory cells were highly reactive to antigen stimulation.

## Discussion

A reference map of commonly recognized epitopes is necessary and important given the association between the size of the functional anti-viral CD8 T cell response and virus control^40–43^. However, recent evidence also point to virus-specific CD8 T cells recognizing common virus infections as bystanders, as well as suspected modulators of pathology, in various disease-contexts classically regarded to not involve virus infections including Alzheimer’s disease^44^ and many human tumors^45^. Here, our reference list of epitopes could be used to elucidate epitope targets and activity of bystander T cells to better understand their disease modulating behavior. Furthermore, bystander CD8 T cells can also serve as therapeutic targets of reactivation as shown by Rosato and colleagues^45^, making our list potentially suited for translational application in cancer immunotherapy as well.

Ranking each epitope according to observed prevalence of reactivity allowed us to further explore phenotypic heterogeneity between antigen-specific populations of CD8 T cells, revealing that non-reactive epitope-specific populations, in addition to senescent Temra-like phenotypes, were associated with CD8^lo^, naïve-like phenotypes, consistent with prior literature from mice that naïve CD8 T cells downregulate CD8 expression and TCR sensitivity through a process of “co-receptor tuning”^18^. Given that the loss of CD8 co-receptor expression was evident and uniform for the majority of observed non-reactive populations, we speculate that the same phenomenon may occur for Temra-like CD8 T cells expressing CD57 and CD45RA while lacking CCR7, CD27 and CD28. These hallmarks, as well as *GZMK* expression, are consistent with replicative senescence^39^. As the infectious history of our donors was unavailable, we cannot know whether these Temra-like cells were actually naïve CD8 T cells or senescent effector CD8 T cells that were primed through infection or vaccination.

Our study highlights two biases related to manually curated epitope databases such as IEDB: (1) prevalence of recognition in actual cohort does not correlate with frequency by which each epitope has been studied, and (2) knowledge on adaptive T cell immunology continues to be biased towards epitopes restricted to HLA-A*02:01. A large number of unrecognized epitopes from common virus infections restricted to non-HLA A*02:01 alleles consequently awaits exploration – e.g. in the context of HAdV-C, B19 and JCPyV virus which had comparable reactivity profiles to FLU-A, EBV and CMV. A broader range of epitopes restricted by more HLA alleles will need to be assessed in future studies in order to understand potential within-host heterogeneity and to enable studies in ethnically diverse populations with low HLA A*02:01 frequency.

## Materials and Methods

### Donor material

Whole blood from 27 anonymous blood donors were obtained at the central blood bank at Rigshospitalet, Copenhagen. PBMCs from 21 hepatitis patients were collected under protocol ID #1999-P004983 “Cell Mediated Immunity in Viral hepatitis”. All patients and healthy donors provided informed consent prior to sampling and according to principles of the Declaration of Helsinki.

### Assembly of DNA-barcode labelled dextran multimer libraries

See supplemental methods for details on epitope selection. Assembly of DNA barcode-labeled dextran multimer libraries were performed using biotinylated DNA-barcodes (LGC Biosearch, 2.17 μM), fluorescent streptavidin-dextran conjugates (Fina Biosolutions Inc., 160 nM), and custom recombinant, biotinylated, UV-cleavable pMHC monomers (50 μg/ml or aprox. 1 μM). Fluorescent dextran conjugates were centrifuged twice at 10,000 g (2 mins, 4°C) to remove aggregates, followed by pre-incubation with biotinylated DNA-barcodes for 30 minutes at 4°C at a molar ratio of 0.5 DNA-barcodes per dextran. Peptide-loaded pMHC monomers were subsequently spun at 3300g, 4°C, 5 minutes, before addition of supernatant to barcoded dextran conjugates at a molar ratio of 16-18 pMHC monomers per dextran. Finally, a premixed freezing buffer was added to a final concentration of PBS, 1.5 μM D-Biotin, 0.1 mg/ml Herring-DNA, 0.5% BSA, 2 mM EDTA, and 5% glycerol. Assembled, DNA barcode-labelled multimers were incubated 20 minutes at 4°C before storage at −21°C. The initial concentration of dextran backbone was 160 nM and the final concentration of assembled multimer was 35.56 nM. See supplemental methods for further details.

### Pooled peptide stimulation

HLA-restricted peptide pools were generated using a liquid handling robot and stored at −21°C until needed. Donor-specific pools were generated on the day of stimulation with 10 μM of each peptide, and used at a final concentration of 1 μM. Cryopreserved PBMCs were thawed using X-Vivo 15 and 5% human serum. All samples were washed twice in 10 mL media before 2-3 million PBMCs were added to PBS-diluted peptides or equimolar DMSO controls.

### Flow cytometry and sorting of multimer-binding CD8 T cells

1.5 μl of each assembled DNA-barcode-labelled multimer was pooled for every stain. Two staining reactions were prepared for each donor. Multimers were first pooled in HLA-restricted pools and subsequently divided into HLA-matching, patient-specific multimer pools. Multimer pools were then concentrated and spun twice to remove aggregates before staining each stimulated/unstimulated sample for 15 minutes at 37°C followed by labelling with surface antibodies and viability dyes at 4°C. See supplemental methods for further details on staining with DNA-barcode labelled multimers as well as combinatorial encoded fluorescent tetramers.

### Amplicon sequencing

Co-attached DNA-barcodes from sorted antigen-specific populations were amplified using *Taq* polymerase and sample-indexed primers as described previously^24^, see supplemental methods for details and supplemental table 3 for PCR primers.

### Sequence analysis

Sequence analysis to calculate significantly enriched DNA-barcodes after cell sorting was run as previously described^24^ using the Barracoda 1.8 web service (https://services.healthtech.dtu.dk/service.php?Barracoda-1.8). See supplemental methods for a detailed description of the Barracoda pipeline.

### Statistical analysis

Flow analysis was done with FlowJo v10.8.1. FlowJo tables and Barracoda outputs were merged and processed for statistical analysis and visualization in R v4.0.5. All correlation plots used Spearman correlations, all two-sample comparisons are performed using Wilcoxon test. Omnibus tests were done using Kruskal-Wallis test, and post-hoc analysis among multiple groups was performed using Dunn’s test with correction for multiple hypothesis testing using Benjamini-Hochberg. All experiments were performed once using multiple technical controls or biological replicates as stated in the figure texts.

## Data Sharing

All data will be made available via email contact to the corresponding author.

## Acknowledgments

We thank technical staff Bente Rotbøll and Anni Flarup Løye for their technical expertise, and all participating donors. We furthermore thank the Memorial Sloan Kettering Cancer Center and Prof. Christina Leslie as well as Dr. Kilian Schober from the University Hospital Erlangen for their wonderful commentary and hospitality in closing this manuscript. The project was funded by the European Research Council, and the ERC StG NextDART and ERC CoG MIMIC; and the Independent Research Fund, Denmark, EliteForsk grant to NPK and SRH.

## Author Contributions

NPK designed the experiments, optimized the assay, generated data, analyzed experimental data, and wrote the manuscript. ED generated experimental data. AKB designed the DNA barcodes, provided consultation and helped designing and performing experiments. UKH maintained and quality-checked the stock of biotinylated DNA barcodes. TT manufactured folded-, biotinylated pMHC monomers. JSK generated data and provided analytical assistance. GN provided bioinformatics assistance. LFV provided experimental assistance. GL provided samples and input to data analysis. SRH is the responsible principal investigator, designed experiments, wrote the manuscript, and provided guidance throughout.

