## Supplemental figures for "Simultaneous analysis of pMHC binding and reactivity unveils virus-specific CD8 T cell immunity to a concise epitope set"

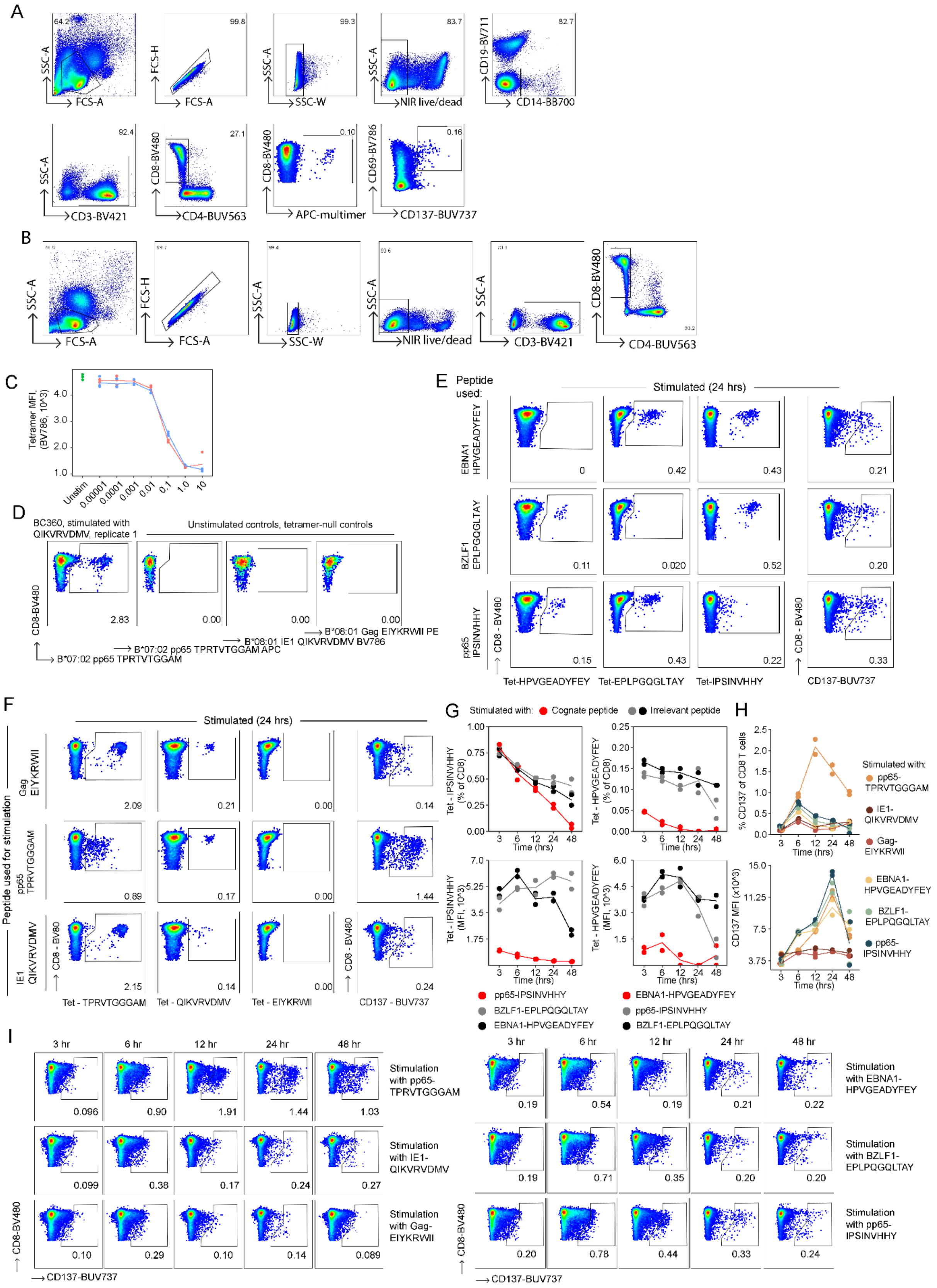

**Supplementary figure 1. Gating controls and additional time series supporting figure 1.**

(A) Gating strategy for pregating CD14-CD19-CD3+CD4-CD8+ T cells.

(B) Gating strategy for pregating CD3+CD4-CD8+ T cells.

(C) BV786 MFI covering tetramer-ELKRKMMYM supporting the limiting dilution experiment covered in Figure 1E.

(D) Gating controls necessary for Figure 1F. A subset of stimulated samples featured a shoulder, which was excluded by gating across all samples.

(E-H) Time series data pertaining to samples stimulated with single peptides presented in Figure 1F. Each sample was subsequently stained using tetramers as indicated on the y-axis. Frequency of tetramer+ cells are measured for bystander (grey and black) and cognate (red) CD8 T cells specific for the stimulated peptide.

(E) Representative flow cytometry gating for time series presented covering HPVGEADYFEY, EPLPGQGLTAY, or IPSINVHHY in donor 373. HPVGEADYFEY, EPLPGQGLTAY, or IPSINVHHY

(F) Representative flow cytometry gating for time series presented covering TPRVTGGAM, QIKVRVDMV, or EIYKRWII in donor 360.

(G) Frequency and MFI of multimer-positive CD8 T cells following stimulation for 3-48 hrs with cognate or irrelevant peptide.

(H) Frequency and MFI of CD137+ CD8 T cells pertaining to the time series conducted in C and D.

(I) Representative CD137 stains following peptide stimulation for 3-48 hrs.

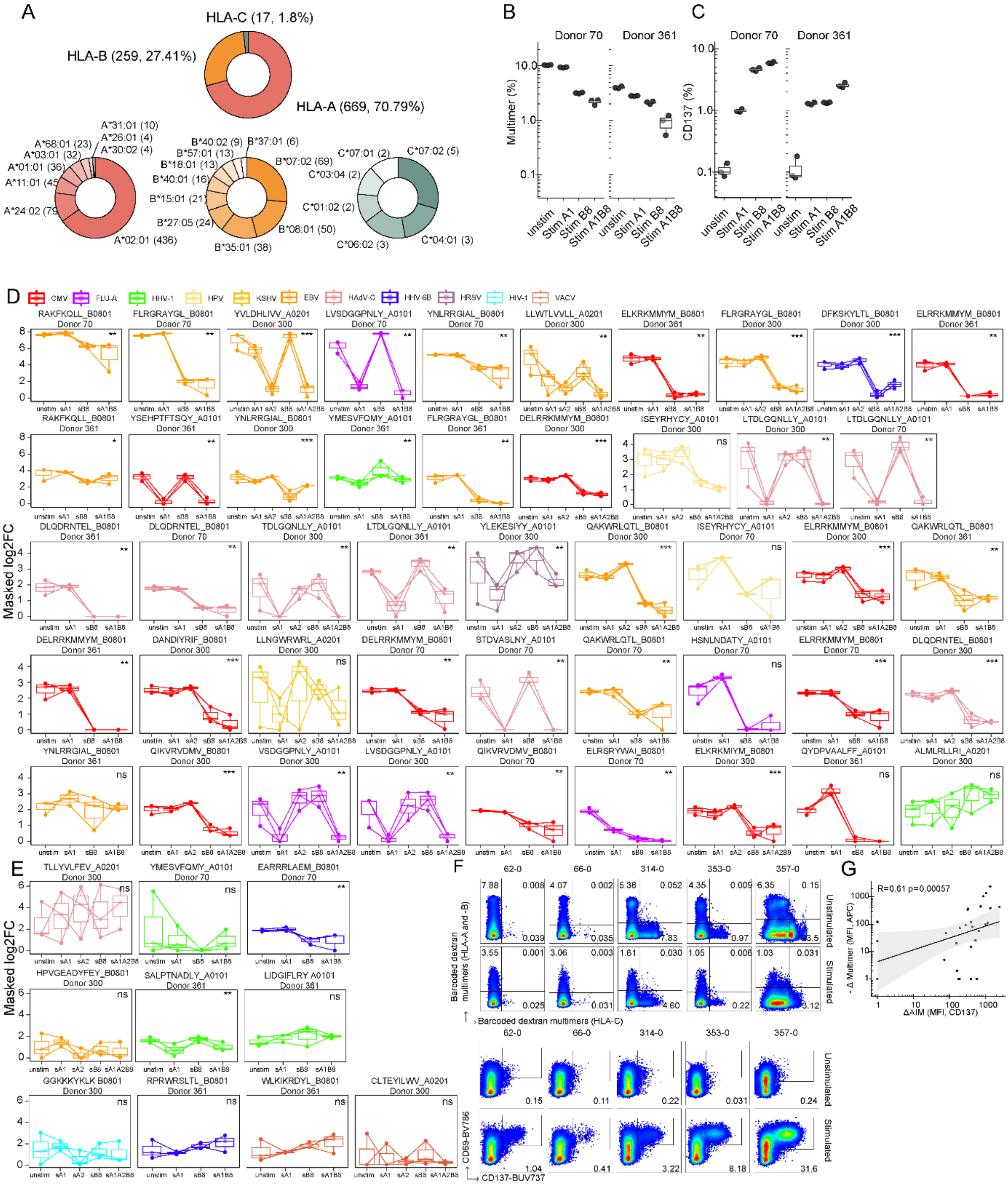

**Supplementary figure 2. Antigen, HLA distribution, and additional flow cytometry data pertaining to figure 2.**

(A) Summary of selected virus epitopes from IEDB per HLA restriction.

(B) Frequency of multimer+ cells among CD8 T cells for triplicate stimulations in addition to Figure 2. Donor 300 was not included as we did not have enough anti-CD137 for all samples.

(C) Frequency of CD137+ cells among CD8 T cells for triplicate stimulations in addition to Figure 2. Donor 300 was not included as we did not have enough anti-CD137 for all samples.

(D-E) Additional virus specific populations identified in triplicate screens for figure 2. unstim, unstimulated. sA1, stimulated with A*01:01-restricted peptides. sA2, stimulated with A*02:01-restricted peptides. sB8, stimulated with B*08:01 restricted peptides. sA1B8 and sA1A2B8 refer to stimulations using recombined pools of peptides. (D) All responses with 2 or more observations across unstimulated triplicates. (E) All responses with only 1 observation across unstimulated triplicates.

(F) Additional representative sorting gates for barcode-labelled multimer stained samples. Single positive gates were sorted into the same tube for PCR amplification and sequencing.

(G) Comparison of increasing -ΔMultimer(MFI) with ΔAIM(MFI, CD137) for samples were only APC-dextran multimers were applied. Samples with negative or zero values were masked to 50% of the lowest observed positive value 7 out of 27 donors/data points were masked this way.

Boxplot bounds show the 25$^{th}$ and 75$^{th}$ percentiles along with the median. Upper and lower whiskers show the range of data unless data points are lower/larger than 1.5xIQR. Correlation and associated p-value was calculated using Spearman's correlation. p-values were calculated using Wilcoxon test in D-E between grouped unstimulated/bystander stimulated samples vs cognate stimulated samples. Dunn's test and corrected for multiple hypothesis testing using Benjamini-Hochberg. *, $p < 0.05$. **, $p < 0.01$. ***, $p < 0.001$. ****, $p < 0.0001$.

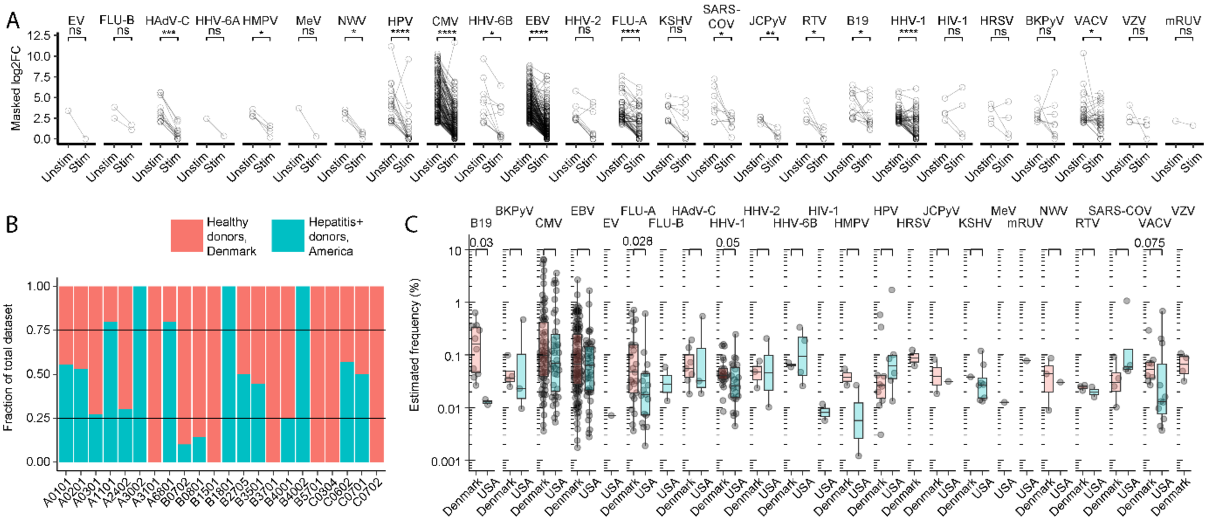


**Supplementary figure 3. Additional gating and comparative analysis of DNA-barcodes.**

(A) Enrichment scores (masked log2FC) for all significantly enriched DNA-barcodes/epitope identities, grouped by virus origin, from both unstimulated controls and stimulated culture conditions. All negative log2FC values were set to 0 (i.e. masked).

(B) HLA distribution across 48 donors from healthy donors (Denmark) and American donors (Hepatitis+). Solid lines mark the 25$^{th}$ and 75$^{th}$ percentile.

(C) Comparison of virus-specific immune responses for HLA alleles with similar distribution (i.e. for HLAs where the median fell within the 25$^{th}$ and 75$^{th}$ percentile in E).

p-values were calculated using a paired Wilcoxon test in A. Boxplot bounds show the 25$^{th}$ and 75$^{th}$ percentiles along with the median. Upper and lower whiskers show the range of data unless data points are lower/larger than 1.5xIQR.

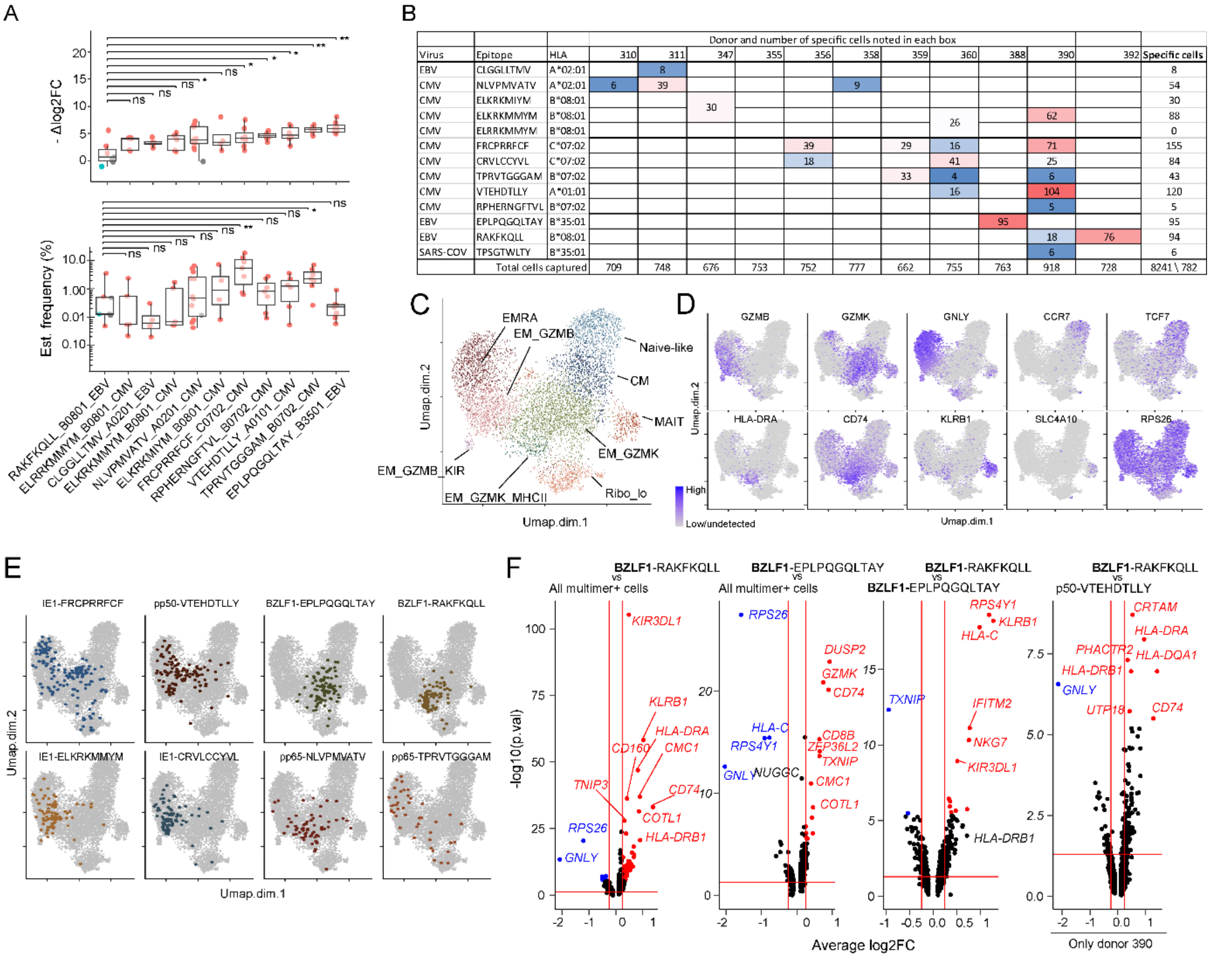


**Supplementary figure 4. Single cell sequencing of virus-specific CD8 T cells using barcode-labelled multimers**

(A) Summary of average reactivity scores and estimated frequencies across our cohort for epitope-specific responses captured using single-cell RNA + TCR sequencing.

(B) Number of cells captured and mapped for each barcoded multimer among 8241 CD8 T cells captured using single-cell RNA + TCR sequencing.

(C) UMAP depicting 9 clusters annotated according differentially expressed markers.

(D) 10 representative markers used for cluster annotation.

(E) Same UMAP as C highlighting the shown antigen-specific CD8 T cells.

(F) Pseudo-bulk differential gene expression using groups annotated as on top of each plot. Right most plot represents the annotated antigen-specificities within a single donor (donor 390). Average log2FC cutoff was set at 0.25. Adjusted p-values were used to color each significantly expressed marker, and the method applied was Bonferroni

**Supplemental Tables**

*These tables are provided as supplemental files*

Table S1. All virus abbreviations

Table S2. Minimal epitope panel and summary statistics

Table S3: DNA-barcodes and amplification primers used.
