## Supplemental methods for "Simultaneous analysis of pMHC binding and reactivity unveils virus-specific CD8 T cell immunity to a concise epitope set"

**Donor material (extended).** PBMCs from Danish blood donors were enriched using Lymphoprep medium and Leucosep tubes for density centrifugation following manufacturer's protocol. PBMCs from 21 hepatitis patients were collected under protocol ID #1999-P004983 "Cell Mediated Immunity in Viral hepatitis". 13 participants had chronic HCV infection, 6 participants had acute HCV infection, 1 participants was spontaneous resolving infection, and 1 participant was positive for HBV. For anonymous blood donors, HLA information were obtained through NGS typing services at DKMS Life Science Lab GbmH (Dresden, Germany). For hepatitis patients, HLA typing was obtained through services at University of Oklahoma Health Sciences Center typing.

**Query of epitopes from common virus infections**. The Immune Epitope Database (IEDB)[23] was queried on October 6th 2020 with the following filters: *Positive assays only, epitope structure: Linear, Organism: Virus (ID: 10239), No B cell assays, No MHC ligand assays, MHC: Restriction Type: Class I, Host: Homo sapiens (human)*, which generated a list of 18.155 epitope entries. We reduced the list to only include 8-15mer epitopes associated with viruses of interest (Supplemental Table 1) and HLA alleles of interest. We reduced the list of entries by excluding structural assays and two model-based assays: *"pathogen burden after challenge"* and "*survival from challenge". We reduced the collected epitope list by only including epitopes with annotated capacity to bind at least one of 25 HLAs of interest excluding serological HLA-typing. Epitopes from human immunodeficiency virus (HIV-1) were included as negative controls. To reduce redundant sequences, epitopes with fewer than 5 assay references in IEDB were removed if they were similar to a sequence with more assay references within the same HLA group. Sequence similarity was here defined as a longest common substring score of 2 amino acids. The final list contained 858 peptide sequences with 945 unique peptide-MHC combinations. 2 µmol lysophilized peptides were purchased at Pepscan (Lelystad, Netherlands) and stored at -21°C upon arrival. Peptides were later rested for 2 hours and dissolved to 10 mM in DMSO and shaken gently for 2 hrs before further alliquoting and storage at -21°C. All peptides (ordered through Pepscan, NL) were controlled by mass-spectrometry analyses. This revealed synthesis or location mistakes related to 16 peptides, which was excluded from analyses: RLNQLESKV(A*02:01), TPRVTGGGA (A*02:01), SAPLPSNRV (A*02:01), QIFLEVQAI (A*02:01), NYLDLSALL (A*24:02), QYDPVAALFF (A*24:02), FYTPLADQF (A*24:02), PYAVCDKCL (A*24:02), QAIRETVEL (A*24:02), RYCCYYCLTL (A*24:02), AYRRRWRRL (A*24:02), SWPDGAELPF (A*24:02), SYAAAQRKL (A*24:02), SYAAAQRKLL (A*24:02), SYKTLREFF (A*24:02), IPVMRKAYL (B*08:01).

**Recombinant expression of UV-cleavable pMHC complexes**. We followed previously published protocols for expression, folding, purification and biotinylation of UV-cleavable pMHC molecules[1,2]. In brief, HLA heavy chains and β2 microglobulin (B2M) were expressed separately in *Escherichia coli* using pET series expression plasmids. Soluble denatured heavy chain and B2M were subsequently isolated as inclusion bodies and refolded in presence of a UV-cleavable peptide followed by subsequent biotinylated (BirA biotin-protein ligase standard reaction kit, Avidity LLC, Auroa, CO) and purified using HPLC (HPLC, Waters corporation, USA). HLA-A*02:01 and HLA-A*24:02 were folded and purified as disulfide-stabilized complexes and stable as empty-loadable monomers as described previously[3]. All MHC-I folded monomers were quality-checked for their concentration, UV degradation, and biotinylation efficiency and stored at −80°C.

**Synthesis and elongation of biotinylated DNA-barcodes.** Biotinylated DNA barcodes were designed, procured and constructed as previously described[4]. However, elongation from single-stranded oligos to double stranded oligos was performed with increased concentrations of DTTs and dNTPs as described here. Two sets of single-stranded oligonucleotides, each with a distinct 25mer "barcode" region described in Xu et al 2019[5], were purchased from LGC Biosearch and delivered in volumes of 100 uM. Oligo set A contained a 5' biotin tag and was joined to the unmodified partially complementary B oligos by annealing and elongation as follows: Oligo combinations were mixed in 384 well plates utilizing 192 wells, and elongated using 5x Sequenase Reaction Buffer Mix (PN 70702, Affymetrix) to a final concentration of 26 µM (Oligo A) and 52 µM (Oligo B). The AxBx combinations were then annealed by heating to 65°C for 2 min and cooled slowly to < 35°C over 15-30 minutes; addition of one reaction Sequenase polymerase (70775Y, Affymetrix), 6.4 mM DTT and 1.28 mM dNTPs, and incubation for 15 minutes at room temperature. The concentration of biotinylated DNA barcodes after elongation was 20.51 µM which was further diluted before aliquoting in PBS+0.5% BSA+1 mg/ml herring DNA+2 mM EDTA to 2.17 µM. Aliquoted plates of elongated DNA barcodes were stored at -21°C. All primers and barcode oligo information are available in Supplementary Table 3.

**Peptide-loading and UV-exchange of UV-cleavable pMHC monomers.** Peptides were diluted from 10 mM to 200 µM in fresh PBS whereas recombinant loadable-pMHC molecules were diluted to 100 ug/ml (aprox. 2 µM). Diluted peptides and loadable-pMHCs[3] were then mixed 1:1 and exposed to UV-light (366 nm) for 1 hr to break cleavable peptides and induce loading of the peptide of interest. Empty-loadable monomers, HLA-A*02:01 and A*24:02, were loaded for 1 hour at 4°C. The final concentration of loaded/exchanged monomers was 50 ug/ml or aprox. 1 µM, and spun at 3300g, 4°C, 5 minutes to pellet aggregates prior to loading.

**Assembly of DNA-barcode labelled dextran multimer libraries.** Assembly of DNA-barcode labelled dextran multimers was performed for each library of HLA-restricted peptide-loaded MHC class I molecules. Streptavidin-conjugated, APC- and PE- dextrans were custom made by Fina Biosolitions Inc. (USA) with the following sets of ratios MOS734: 4.2 APC/Dex, 4.9 SA/Dex and MOS722: 5.4 PE/Dex, 6.2 SA/Dex. Fluorescent dextrans were first spun at 10.000g, 4°C, 2 minutes and transferred to fresh Eppendorf tubes to remove pellets and repeated a second time. Fluorescent dextrans were then preincubated with 0.5 DNA-barcodes per dextran for 30 minutes at 4°C followed by addition of 16-18 pMHC of interest per dextran and incubation for 30 minutes at 4°C. Finally, a premixed freezing buffer was added to a final concentration of PBS, 1.5 µM D-Biotin, 0.1 mg/ml Herring-DNA, 0.5% BSA, 2 mM EDTA, and 5% glycerol. Assembled, DNA barcode-labelled multimers were then incubated for at least 20 minutes at 4°C before storage at -21°C until needed. The initial concentration of dextran backbone was 160 nM and the final concentration of assembled multimer was 35.56 nM.

**Peptide pool stimulation (extended)**. Dissolved peptides were pooled using a liquid handling robot by taking 5 µL of each well and depositing in a common reservoir. Total volume was then measured and split 5-ways into labelled Eppendorf tubes and stored at -21°C. On the day of stimulation peptide pools were thawed and recombined into donor-specific pools by adding X*0.25 µL, where X corresponds to the number of HLA-restricted peptides in the HLA-restricted pool. The donor-specific pool of peptides  were then diluted to 250 µL. Due to the high number of A*02:01-restricted epitopes A*02:01 positive donors were stimulated with two separate A*02:01-restricted peptide pools, with approximately  220 peptides each, along with a separate non-A*02:01-restricted pool to avoid lethal dosages of DMSO. Each pool contained 10 µM of each peptide, and used at a final concentration of 1 µM after addition of donor PBMCs.

**Flow cytometry and sorting of multimer-binding CD8 T cells (extended).** Concentration of multimer panels was performed using Vivospin 8 columns to 60 µl volume per stain. Donor-specific multimer pools were spun twice to remove aggregates at 10.000g, 2 min, 4°C transferring the pool to new Eppendorf tubes. Volume of the aggregate-reduced multimer pool was measured, and adjusted by adding barcode-cytometry buffer (PBS + 0.5% BSA + 100 µg/mL herring DNA + 2 mM EDTA). A 5 µL aliquot of the aggregate-reduced multimer pool was then stored at -21°C for later PCR amplification. Cells were spun gently at 390g, 5 min, 4°C and followed by resuspension in barcode-cytometry buffer and washed at 390g. Pellets were the resuspended in ~60 µL multimer pools supplemented with 1 µM Dasatinib and incubated for 15 minutes at 37°C before staining with surface antibodies/viability-dye and cell sorting. Sorting gates were set individually for each donor, but constant between stimulation conditions. CD14-CD19-CD3+CD4-CD8+ multimer-binding cells were sorted using BD FACSAria Fusion. Surface-antibodies and near-infrared fixable-viability dye were pooled in concentrations giving by our key resource table in the supplemental methods. Cells were incubated 30 minutes at 4°C, and washed three times at 390°C before fixation and cell sorting. Alternatively, cells were fixed overnight at 4°C with 1% PFA. All washing steps as well as the sorting buffer (50 µl) was performed in BCB. Reservoirs and Eppendorf tubes for sorting were saturated with 2% BSA prior to staining. For intracellular staining and immunophenotyping tetramers were spun at 3300g, 5 mins, 4°C to remove pellet. This was repeated twice before staining at double the concentration descried previously[6].

**Amplicon sequencing (extended).** PCR machines were setup with the following cycles 95 °C 10 min; 36 cycles: 95 °C 30 s, 60 °C 45 s, 72 °C 30 s; and 72 °C 4 min. Amplification was also performed on non-template controls, and on three samples of the original, donor-specific multimer pool (diluted 1:50.000). The later is necessary to estimate the baseline distribution of DNA-barcodes / multimers from which a signal on sorted cells can be enriched. Pooled products were purified using QIAquick PCR purification kit. 200 ng cDNA were diluted in 25 µl nuclease-free water and send for Ion Torrent PGM™ sequencing at Primbio Inc. (USA). 3-6 million reads per chip were ordered depending on the number of pooled samples with an expected average of 150 high-quality reads per barcode.

**Sequence analysis (extended)**. In brief, Barracoda calculates log fold change (log2FC) and p-values using EdgeR[34] comparing sorted products to the mean of triplicate baseline values from amplified multimer pools originating before staining. In our application, the barcode-wise log2FC ($M_{gk}$) is calculated as follows[35]:

$$M_{gk} = log2(\frac{Y_{gk(sorted\ population)}/N_{k(sorted\ population)}}{Y_{gk(baseline)}/N_{k(baseline)}})$$

$Y_{gk}$ is the number of clonally reduced reads (UMIs) counted for each DNA barcode $g$ in sample $k$. $N_k$ represents the total number of UMIs counted for the given sample. Triplicate mean baseline values for each DNA barcode is used as input for the denominator. P-values were corrected using the Benjamini-Hochberg procedure. Specific barcodes with FDR < 0.1% were defined as significant. All significant hits needed to represent at least 1/1000 reads to be defined as significant: A threshold used to exclude hits due to low coverage in input samples. Estimated frequency of a given population of significantly enriched, epitope-specific CD8 T cells were calculated as follows[24]:

$$Est.freq_i = pMHC\ multimer(\%)_k \cdot Count.fraction_i$$

$pMHC\ multimer(\%)_k$ represents the frequency of multimer-specific CD8 T cells observed during cell-sorting of a given sample on $k$ dextran fluorochrome whereas $Count.fraction_i$ represents the given count fraction of the multimer $i$ of interest.

**Single cell analyses of pMHC multimer binding CD8 T cells**. Cryopreserved PBMCs stained using barcoded pMHC multimers (as described earlier) and standard CD8 T cell identification markers. Furthermore, cells were treated with 2.5 µl of Human TruStain FcX™ Fc Blocking reagent (10 min at 4 °C) in a total of 50 µl cell suspension. 0.5 µl of a unique hashing antibody (Biolegend, TotalSeq™-C0251 anti-human Hashtag 1-10 Antibodies) was added to a given sample, followed by incubation for 15 min at 4 °C. pMHC multimer positive T cells were sorted and mixed across samples. ~15,000 cells were loaded (based on 55% recovery from 27,500 sorted cells) onto a Chromium Controller to yield a maximum of 9,000 cells with an intermediate/high doublet rate (6,9%). We utilize the 10x Genomics 5' v2 chemistry that allows the cell barcode to be appended at the 5'-end of transcripts, which is for capturing all V(D)J-, pMHC-, and hashing-associated associated barcodes as described previously[7]. The downstream processing were conducted according to manufacturer's instruction (10x Genomics), and the different products (GEXs, TCRs and barcodes) were sequenced on a NovaSeq running a 150 paired-end program.

Gene expression, hashing-associated reads and pMHC-associated reads were processed using Cellranger multi (10X genomics) with hashing antibody and pMHC annotations. The relevant outputs were merged into a single Seurat object followed by removal of doublets using HTOdemux (Seurat) and assembly of consensus TCR sequences for each cell. pMHC-associated reads were demultiplexed and antigen-specific clonotypes manually called using donor-specific heatmaps. Low-quality cells were removed prior to clustering and differential gene expression analysis. Only cells showing less than 10% mitochondrial gene expression and more than 100 expressed genes were included. Gene expression was transformed using default settings of SCTransform followed by clustering using

FindClusters(resolution = 0.6) and FindClusters(resolution = 0.4) to identify two minor outlier clusters ("MAIT" and "EM_GZMB_KIR"). Number of principal components used for clustering was set to include 90% cumulative variation. Mitochondiral genes, TCR genes (e.g. TRAV) and ribosomal genes were excluded from the list of variable features prior to clustering and dimensionality reduction using umap.

**Immunophenotyping using combinatorial encoded tetramers.** Tetramers were generated and used as described previously[6,8]. In brief, a surplus of 100 µg/ml pMHC (~ 2 µM[1]) were mixed into 100 µg/ml (~0.55 µM) streptavidin at 30 molar ratio followed by 30 minutes incubation at 4$^{o}$C and addition of 1 µL 2% FCS buffer per 9 µL construct. Streptavidin color combinations were premixed at equal molar ratios. All fluorescent streptavidins were centrifuged at 10.000 g, 2 minutes, 4$^{o}$C to remove aggregates before assembly with folded pMHC. Tetramers were spun three times at 3300g, 5 minutes at 4*C and transferred to empty wells of a 384 well plate in-between centrifugations. Tetramers were subsequently added one at a time to 50 µl cells resuspended in 2% FCS (FACS buffer). Staining concentration was 1.5 µl per 50 µl cells.

**Computational analysis of flow cytometry data**. Number tetramer+ cells were evaluated using flowJo and compensated fluorescence intensities were exported for each tetramer+ population with more than 20 tetramer+ cells. Maximum 5000 cells were exported from each tetramer+ population. 5000 cells were also exported from each population of bulk CD8 T cells. The R package CATALYST[9,10] and flowcore[11] were used for processing before annotation and conversion to a SingleCellExperiment object where MDS, UMAP and flowSOM analysis was performed. CD45RA, CCR7, CD95, CD127, GzmB, CD57, Ki67, CD27, CD28 and GzmB were selected for clustering. CD25, CD4, CD3, CD8, tetramer-dedicated channels, CD14, NIR and CD19 were deselected. 8 clusters were chosen for flowSOM clustering, as an elbow plot featured loss of new information at 8 clusters along with suitable resolution of memory, effector and naïve subsets.

**Key ressources**

| Antibodies, dyes, and fluorescent streptavidin (SA) | SOURCE | IDENTIFIER |
|---|---|---|
| Anti-CD3 BUV395, clone: UCTH1, staining conc: 1 / 100 uL | BD | Cat# 563546 |
| Anti-CD4 BUV596, clone: SK3, staining conc: 0.5 / 100 uL | BD | Cat# 612912 |
| Anti-CD137 BUV737, clone: 4B4-1, staining conc: 2.5 / 100 µL | BD | Cat# 741861 |
| Anti-CD3 BV421, Clone: SP34-2, staining conc: 2 / 100 µL | BD | Cat# 562877 |
| Anti-CD8 BV480, Clone: RPA-T8, staining conc: 1 / 100 µL | BD | Cat# 566121 |
| Anti-CD19 BV711, Clone: SJ25C1, staining conc: 2.5 / 100 µL | BD | Cat# 563036 |
| Anti-CD14 BB700, Clone: MφP9, staining conc: 0.2 / 100 µL | BD | Cat# 566465 |
| Anti-CD69, BV786, Clone: FN50, staining conc: 2.5 / 100 µL | BD | Cat# 563834 |
| Anti-CD19 FITC, Clone: 4G7, staining conc 6.25 / 100 µL | BD | Cat# 345776 |
| Anti-CD14 FITC, Clone: MφP9, staining conc: 3.12 / 100 µL | BD | Cat# 345784 |
| Anti-CD95 BUV395, Clone: DX2, staining conc: 2.5 / 100 ul | BD | Cat# 740306 |
| Anti-CD28 BUV737, Clone: CD28.2, staining conc: 1.25 / 100 ul | BD | Cat #612815 |
| Anti-CD27 BV605, Clone: O323, staining conc: 2.5 / 100 ul | Biolegend | Cat #302830 |
| Anti-ki67 BV650, Clone: B56, staining conc: 1.25 / 100 ul | BD | Cat #563757 |

| | | |
|---|---|---|
| Anti-CD45RA BV711, Clone: HI100, staining conc: 2.5 / 100 ul | BD | Cat #563733 |
| Anti-CD57 BV785, Clone: QA17A04, staining conc: 1 / 100 ul | Biolegend | Cat #393329 |
| Anti-CCR7, FITC, Clone: G043H7, staining conc: 5 / 100 ul | Biolegend | Cat #352116 |
| Anti-CD25, BB700, Clone: M-A251, staining conc: 2.5 / 100 ul | BD | Cat #566447 |
| Anti- GzmB, PE-CF594, Clone: GB11, staining conc: 1.25 / 100 ul | BD | Cat #562462 |
| Anti-CD127, APC-R700, Clone: HiL-7R-M21, conc: 5 / 100 ul | BD | Cat #565185 |
| Anti-CD14, APC-Cy7, Clone: MφP9, staining conc: 2.5 / 100 µL | BD | Cat #557831 |
| Anti-CD19, APC-Cy7, Clone: SJ25C1, staining conc: 2.5 / 100 µL | BD | Cat #557791 |
| Anti-CD4, FITC, Clone: sk4, staining conc: 1.2 / 100 ul | BD | Cat #345768 |
| Anti-CD14, FITC, Clone: MφP9, staining conc: 3.125/ 100 ul | BD | Cat #345784 |
| Anti-CD19, FITC, Clone: 4G7, staining conc: 6.25 / 100 ul | BD | Cat #345776 |
| Anti-CD40, FITC, Clone: LOB7/6 , staining conc: 2.5 / 100 ul | Bio-Rad | Cat # MCA1590F |
| LIVE/DEAD™ Fixable Near-IR Dead Cell Stain Kit | Thermo Fischer | Cat# L34976 |
| Anti-PD-1, BV421, Clone: EH12.1, staining conc: 2.5 / 100 ul | Biolegend | Cat #329920 |
| Anti-CD57, PE-Cy7, Clone: QA17A6, staining conc: 2.5 / 100 ul | Biolegend | Cat #393310 |
| Anti-eomes, BUV395, Clone: X4-83, staining conc: 2.5 / 100 ul | BD | Cat #567171 |
| Anti-tbet, BV786, Clone: O4-46, staining conc: 2.5 / 100 ul | BD | Cat #564141 |
| Anti-CD137, BV605, Clone: HIL-7R-M21, conc: 1.25 / 100 ul | BD | Cat #562662 |
| Anti-CD38, BUV737, Clone: HB7, staining conc: 0.625 / 100 ul | BD | Cat #612824 |
| Anti-CD3, R718, Clone: UCHT1, staining conc: 0.9 / 100 ul. | BD | Cat #566953 |
| Anti- KIR2DL1/S1/S3/S5, BB700, Clone: HP-MA4, staining conc: 5 / 100 ul | BD | Cat #752515 |
| SA-BV786 | BD | Cat# 563858 |
| SA-PE | Biolegend | Cat# 405204 |
| SA-PE-Cy7 | Biolegend | Cat# 405206 |
| SA-APC | Biolegend | Cat# 405243 |

| | | |
|---|---|---|
| TotalSeq™-C0251-C0260 anti-human Hashtag 1-10 | Biolegend | Cat# 394661, 394663, 394665, 394667, 394669, 394671, 394673, 394675, 394677, 394679 |
| Custom buffers: | | |
| Custom buffer: Barcode-freezing buffer | N/A | PBS, 1.5 µM D-Biotin, 0.1 mg/ml Herring-DNA, 0.5% BSA, 2 mM EDTA, 5% glycerol |
| Custom buffer: Barcode-cutometry buffer | N/A | PBS, 0.1 mg/ml Herring-DNA, 0.5% BSA, 2 mM EDTA. |
| Biological samples | | |
| Whole blood samples and derivatives including peripheral blood mononuclear cells (PBMCs)) from anonymous blood donors | Copenhagen University Hospital (Region H) | N/A |
| PBMCs from chronic and acute hepatitis C and B patients | Massachusetts General Hospital | N/A |
| Kits, chemicals, peptides, and recombinant proteins | | |
| Taq PCR Master Mix Kit | Qiagen | Cat# 201445 |
| Male AB human serum | Sigma Aldrich | Cat# H3667 |
| X-Vivo 15 | Lonza Bioscience | Cat# BE02-060Q |
| Fetal Bovine Serum (FBS) | Gibco | Cat# 10500-064 |
| DMSO, sterile. | Sigma Aldrich | Cat# 276855-100mL |
| Lymphoprep™ | Stem cell | Cat# 07861 |
| Leucosep™ tubes | Greiner | Cat# 227290 |
| E-gel Double comb Agarose Gels, 2% | Invitrogen | Cat# G601802 |
| Brilliant stain buffer plus | BD | Cat# 566385 |
| QIAquick spin columns | Qiagen | Cat# 28115 |
| Recombinant human UV-cleavable pMHC | Prepared in lab | N/A |
| Recombinant empty-stabilized pMHC | Prepared in lab | N/A |

| | | |
|---|---|---|
| Sequenase Reaction Buffer mix | Affymetrix | Cat# PN 70702 |
| Sequenase polymerase | Affymetrix | Cat# 70775Y |
| DTT | Thermo Fischer | Cat# R0861 |
| dNTP | Life Technologies | Cat# 18427-088 |
| Tween 20 | Sigma Aldrich | Cat# P0379-100ml |
| 2 umol, crude peptides. | Pepscan | Custom order |
| D-biotin | Avidity | Cat# Bio200 |
| Glycerol | Sigma Aldrich | Cat# 1370281000 |
| Bovine Serum Albumin (BSA) | Sigma Aldrich | Cat# A7906-100g |
| Vivaspin 6 columns, 100.000 Dalton | Sartorius | Cat# VS0642 |
| EDTA | Sigma Aldrich | Cat# 03690-100mL |
| Dasatinib | Sigma Aldrich | Cat # CDS023389 |
| Nuclease-free water | Fischer Scientific | Cat# AM9937 |
| Autoclaved PBS | Prepared in bulk | N/A |
| herring DNA | Sigma Aldrich | Cat# D3159-10g |
| PE-labelled SA-Dextran (PE/SA/Dextran, 5.4:5.4:1) | Fina Biosolutions | Custom order |
| APC-labelled SA-Dextran (APC/SA/Dextran, 4.2:4.5:1) | Fina Biosolutions | Custom order |
| QIAquick PCR purification kit | Qiagen | Cat# 28104 |
| FoxP3 / Transcription factor staining buffer set | eBioscience | Cat# 00-5523-00 |
| Human TruStain FcX™ | Biolegend | Cat# 4422302 |
| SPRIselect beads | Bechman Coulter | Cat #B23318 |
| 50 bp DNA ladder | Thermo Fischer | Cat# 10416-014 |
| Software and algorithms | | |
| FlowJo V10.8.1 | BD | |
| Barracoda | Public webservice | https://services.healthtech.dtu.dk/service.php?Barracoda-1.8 |

| | | |
|---|---|---|
| R 4.2.3 | | https://cran.r-project.org/bin/windows/base/old/4.0.0/ |
| R Package: Tidyverse | [12] | https://cran.r-project.org/web/packages/tidyverse/index.html |
| R Package: ggpubr | Alboukadel Kassambara | https://cran.r-project.org/web/packages/ggpubr/index.html |
| R Package: cowplot | Wilke lab, The University of Texas | https://wilkelab.org/cowplot/articles/introduction.html |
| R Package: MetBrewer | Blake Robert Mills https://www.blakerobertmills.com/ | https://github.com/BlakeRMills/MetBrewer |
| R Package: ggbeeswarm | Erik Clarke, Scott Sherrill-Mix, and Charlotte Dawson | https://github.com/eclarke/ggbeeswarm |
| R package: Seurat | Satija Lab, New York Genome Center | https://satijalab.org/seurat/ |
| R Package: xlsx | Cole Arendt | https://cran.r-project.org/web/packages/xlsx/index.html |
| Oligonucleotides | | |
| Forward primer with varying SampleIDs and iontorrent adapters | LGC Biosearch | Table S3 |
| Reverse primer with iontorrent adaptors | LGC Biosearch | Table S3 |
| Biotinylated single-stranded DNA (Oligo Ax). | LGC Biosearch | Table S3 |
| Single-stranded DNA By. | LGC Biosearch | Table S3 |
